# Wdr62 and Patronin cooperate to organize distinct acentrosomal microtubule networks during *Drosophila* oogenesis

**DOI:** 10.64898/2026.04.19.719433

**Authors:** Jéssica Cabrita, Ana Ferreira-Silva, Telmo Pereira, Ana Pimenta-Marques

## Abstract

Acentrosomal microtubule (MT) organization is essential in many differentiated cells, yet the mechanisms that generate and maintain these networks remain poorly understood. Here, we identify Wdr62 as a conserved regulator of acentrosomal MT organization during *Drosophila* oogenesis. Wdr62 localizes to cortical non-centrosomal microtubule-organizing centers (ncMTOCs) and is required for proper localization of the conserved components Shot and Patronin. Loss of Wdr62 disrupts MT organization and oocyte polarity and reduces MT growth events during mid-oogenesis. Genetic analyses further reveal that Wdr62 functionally cooperates with Patronin and the MT-severing enzyme Katanin 80 to organize the acentrosomal MT network. Our data support a model in which Wdr62 promotes stabilization of MT minus ends through Patronin while Katanin-mediated severing generates new MT seeds that amplify the MT network. Moreover, Wdr62 and Patronin function together during assembly of the acentrosomal meiotic spindle, and their loss compromises fertility. Given that WDR62 mutations are associated with human disease in cell types that rely on acentrosomal MT organization, such as neurons, these findings provide insight into how disruption of WDR62-dependent MT organization may contribute to disease beyond female infertility.

## INTRODUCTION

Tight regulation of the microtubule (MT) cytoskeleton is essential for key cellular functions, including cell shape, polarity, migration, and division^1,2^. The geometry of MT networks critically depends on microtubule organizing centers (MTOCs), which direct the polymerization, stabilization, and anchoring of MTs^1–3^. In parallel, microtubule-associated proteins (MAPs) modulate MT dynamics and organization, thereby shaping the architecture and functional properties of the MT cytoskeleton^2^.

The centrosome is the best characterized MTOC, critical for generating radial MT arrays that regulate cell shape and polarity during interphase, and for assembling the mitotic spindle during cell division^4–6^. However, most cells in multicellular organisms are not actively dividing. Many exit the cell cycle or undergo terminal differentiation, and in these contexts centrosomal MTOC activity is generally attenuated^1,3,7^, such as in neurons and epithelial cells, or, in extreme cases, these structures can be completely lost, such as in oocytes^8–10^. Consequently, MT organization in these cells relies on acentrosomal pathways. These pathways include the generation of MTs from other cellular sites, termed non-centrosomal MTOCs (ncMTOCs). These sites can occupy large surfaces, such as the apical surface of epithelial cells, or restricted cortical regions, such as in mammalian karyosomes^11–13^ and *Drosophila* oocytes^14^. In contrast to centrosome-mediated MT organization, which has been extensively characterized, the mechanisms underlying acentrosomal MT organization remain comparatively less understood.

Understanding how acentrosomal MT networks are assembled and organized is particularly important in polarized cells, such as oocytes, neurons, and epithelial cells, where they play essential roles in intracellular transport, cell polarity and tissue architecture^15–17^. In *Drosophila* oocytes the acentrosomal MT network is essential for development, as it underlies the transport and spatial organization of key determinants required for embryonic patterning^14,18,19^. The *Drosophila* oocyte provides a powerful model to study acentrosomal MT organization. During mid-oogenesis, centrosomes are inactivated as MTOCs^14^, yet a highly polarized MT network is established, to which ncMTOCs localized along the antero-lateral cortex contribute^14^. Previous work showed that oocyte ncMTOCs contain the conserved MT minus-end protein Patronin^14^ (CAMSAP in mammals), which stabilizes minus-ends of non-centrosomal MTs in different cell types^20–22^. Patronin forms a complex with the sole *Drosophila* spectraplakin, Short Stop (Shot), which anchors the complex to the cortical actin cytoskeleton. From these ncMTOCs, MTs grow toward the posterior of the oocyte, establishing overall cytoskeletal polarity^14,23,24^. However, how MT growth is generated and organized from ncMTOCs to generate a dense, dynamic, and polarized network throughout the oocyte cytoplasm remains unclear. Canonical MT nucleation mediated by γ-tubulin has been excluded as a contributor to MT growth in the oocyte, based on its absence from cortical regions^14^. In addition to MT generation at defined MTOCs, MT networks can also be amplified through severing-based mechanisms, in which enzymes such as Katanin sever MTs, generating new MT fragments that serve as templates for further growth^25,26^. Since Patronin co-immunoprecipitates with the MT-severing enzyme Katanin 80 (Kat80) from *Drosophila* ovary extracts, it has been proposed that MTs may arise through severing and rearrangement of pre-existing MTs mediated by Kat80^14^. Nevertheless, the mechanisms that coordinate MT growth, minus-end stabilization, and severing to sustain and amplify acentrosomal MT networks remain poorly understood in oocytes and in other cellular contexts, such as neurons, where severing-based mechanisms contributes to the assembly of MT arrays^25,26^.

Wdr62 is a conserved WD-repeat protein 62 best known for its role in mitotic spindle organization^27,28^ and neural progenitor proliferation^29–32^, and mutations in WDR62 are the second most common cause of primary microcephaly in humans^33^. Recent work has also implicated Wdr62 in non-centrosomal MT organization in *C. elegans* intestinal cells^34^, raising the possibility that it contributes to the coordination of acentrosomal MT growth and organization across developmental contexts. Here, we investigated whether Wdr62 contributes to the assembly and organization of acentrosomal MT networks during *Drosophila* oogenesis. We show that Wdr62 localizes to cortical ncMTOCs and is required for proper recruitment of the conserved ncMTOC components Shot and Patronin to the antero-lateral cortex of the oocyte. Consistent with this, Wdr62 is required for the polarized localization of axis-determining proteins, a process dependent on ncMTOC-mediated MT organization. We further demonstrate that Wdr62 promotes persistent MT growth and cooperates with the conserved MT-severing enzyme Kat80 to amplify MT growth events during mid-oogenesis. Finally, Wdr62 and Patronin function together during acentrosomal meiotic spindle assembly and are required for female fertility. Together, our findings reveal how conserved ncMTOC components are redeployed to organize distinct acentrosomal MT networks during different stages of oogenesis.

## RESULTS

### WDR-62 localizes to the antero-lateral cortex and colocalizes with ncMTOC components

Wdr62 has been extensively studied at centrosomes, where it has been implicated in mitotic spindle assembly and the maintenance of spindle pole integrity^27–31,35–37^. However, previous work in *C. elegans* showed that Wdr62 does not localize to centrosomes but instead associates with sites of acentrosomal MT growth in different epithelia and neurons. In intestinal epithelial cells, Wdr62 functions as an ncMTOC component and is required for proper MT organization^34^. Its conservation in organisms lacking centrosomes further suggests an evolutionarily conserved role in acentrosomal MT organization^34^.

To determine whether Wdr62 contributes to acentrosomal MT organization in the *Drosophila* oocyte, we first examined its localization throughout oogenesis. In *Drosophila*, oogenesis begins with the formation of a 16-cell cyst, in which one cell is specified as the oocyte while the remaining cells differentiate into nurse cells (**Fig. 1a**). Early in oogenesis, centrosomes from the nurse cells migrate into the oocyte and cluster at its posterior, forming a large MTOC from which MTs extend. At mid-oogenesis, centrosomes are inactivated as MTOC^38^, and MTs grow from ncMTOCs that localize to the antero-lateral cortex and are composed by Shot and Patronin^14^ (**Fig. 1a**). These ncMTOCs support MT growth, thereby establishing a polarized MT network with minus ends at the cortex and plus ends oriented toward the posterior^14,19,39,40^. A protein trap line tagging all *Wdr62* isoforms was used to visualize endogenous Wdr62 protein^41^. During early stages of oogenesis (germarium to stage 4), when centrosomes are active as MTOCs, Wdr62::GFP did not show obvious localization at centrosomes (**Fig. 1b, c)**. Quantification of colocalization between Wdr62::GFP and the centrosomal marker D-plp revealed detectable overlap in fewer than 20% germarium and in only 8% of stage 4 oocytes, indicating that Wdr62 is rarely associated with centrosomes during oogenesis (**Fig. 1b-d**). Consistent with this, in the germarium, Wdr62::GFP appeared enriched in regions overlapping with Shot localization, which is known to associate with the fusome, an actin– and spectrin-rich structure that plays a critical role in establishing MT polarity during early oogenesis^42,43^.

**Figure 1.**
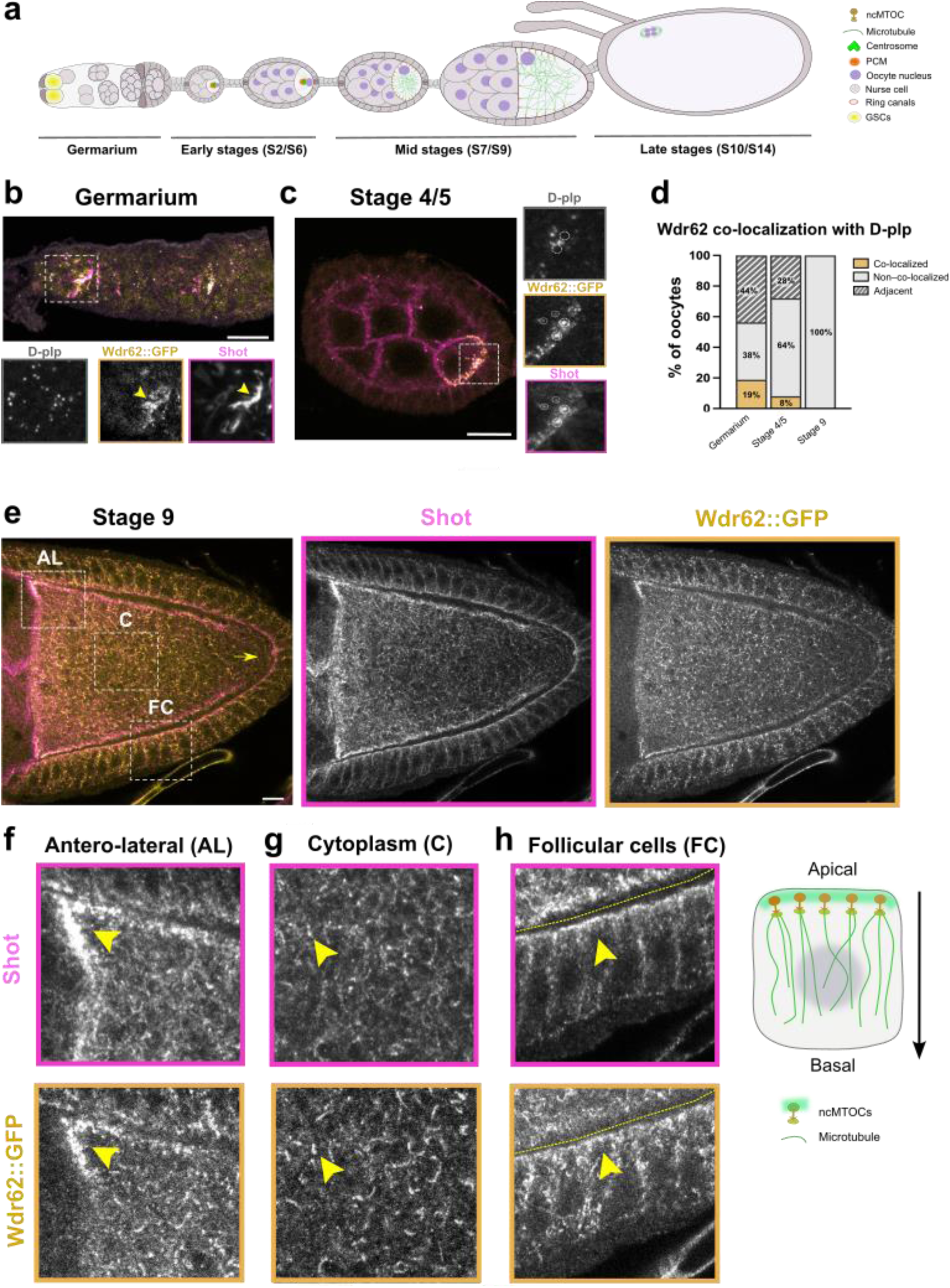
– Wdr62 localizes to antero-lateral cortical ncMTOCs and MT-associated regions during oogenesis. **a**) Schematic of *Drosophila* oogenesis. Oogenesis begins with the asymmetric division of a germ line stem cell (GSC) generating a cystoblast that undergoes four successive mitotic cycles with incomplete cytokinesis, forming a cyst of 16 cells interconnected through intercellular cytoplasmic bridges. One cell adopts oocyte fate, while the remaining cells become nurse cells. Egg chambers bud from the germarium and mature as they move posteriorly. Early in oogenesis most centrioles from each of the nurse cells migrate into the oocyte and cluster together, forming a very large MTOC that organizes the MT cytoskeleton from stages 2 to 6. Around stage 7 centrosomes are inactivated as MTOCs and ncMTOCs are assembled at the antero-lateral cortex. The oocyte remains arrested in meiotic prophase I until stage 13. At this stage, meiosis resumes, marked by nuclear envelope breakdown (NEBD) and the assembly of the meiotic spindle during stages 13–14. Then meiosis arrests again in metaphase I and remains in this state until the egg is activated upon passing through the oviduct. **b,c**) Representative images of Wdr62::GFP localization during early stages of oogenesis. Insets show magnified views of boxed regions. **b**) In the germarium, Wdr62::GFP (yellow) predominantly co-localizes with Shot (magenta) at the fusome, as defined by its characteristic morphology and known Shot localization (arrowhead). **c**) In stage 4/5 egg chambers, Wdr62::GFP co-localizes with Shot at the posterior region of the oocyte (dashed circles) but not with D-plp. **d**) Quantification of oocyte co-localization profile between Wdr62::GFP and the centriole marker D-plp (white). N = 16 (germarium), 20 (stage 4/5) and 28 (stage 9), respectively. **e**) Representative images of stage 9 oocytes showing the localization of Wdr62::GFP and Shot. Wdr62::GFP co-localizes with Shot at the antero-lateral cortex and like Shot, is excluded from the posterior cortex (arrow). In the cytoplasm, both proteins display a filamentous distribution: Note that Wdr62::GFP localizes on curved filaments. Dashed boxes correspond to magnified views in **f-h**. **f-h**) Wdr62::GFP is enriched at the anterior-lateral (AL) region of the oocyte cortex **(f**); within the oocyte cytoplasm (C) **(g);** and at the apical region of the surrounding follicular epithelial cells (FC) (**h**), Arrowheads indicate regions where Shot and Wdr62::GFP colocalize. More than 15 oocytes were analyzed per stage. Scale bars, 10 μm.

Localization of Wdr62 was next examined during later stages of oogenesis, when centrosomes are attenuated as MTOCs^38^ and ncMTOCs are established at the antero-lateral cortex of the oocyte^14^. At stage 9, Wdr62::GFP appeared enriched at the antero-lateral regions of the oocyte cortex (**Fig. 1e, f**). This localization pattern closely resembled that of the known ncMTOC components Shot and Patronin^14^. Consistent with this, Wdr62::GFP colocalized with both Shot (**Fig.1e**) and Patronin (**Supplementary Fig. 1**) at the antero-lateral cortex. Like Shot and Patronin, Wdr62 is excluded from the posterior cortex of the oocyte (**Fig. 1e, arrow**). In addition to its cortical enrichment, Wdr62::GFP was also detected throughout the oocyte cytoplasm. Both Wdr62 and Shot display a mesh-like distribution, with a gradient characterized by a denser filamentous network toward the anterior and a sparser distribution toward the posterior of the oocyte (**Fig. 1e, g**). This pattern resembles the previously described cytoplasmic actin mesh in the oocyte^44^, and is in line with the reported actin-MT crosslinking activity of Shot in other contexts^45,46^.

In addition to its localization in the germline, Wdr62::GFP was detected along MTs in the follicular epithelium (**Fig. 1e, h**) and enriched at the apical surface (**Supplementary Fig. 1b**), where Shot-containing ncMTOCs organize apical–basal MT arrays^14,47^ (**Fig. 1h**).

Together, these results show that Wdr62 is not prominently associated with centrosomes during oogenesis but instead localizes to cellular regions enriched in MTs or ncMTOCs. Such regions include the fusome, the oocyte cytoplasm during mid-oogenesis, and ncMTOCs at the oocyte cortex and the apical surface of follicular epithelial cells. These observations support a role for Wdr62 in acentrosomal MT organization during oogenesis.

### WDR-62 is required for proper organization and polarity of the oocyte microtubule network

Given its localization, we next asked whether Wdr62 is required for proper organization of the oocyte MT cytoskeleton during mid-oogenesis, when centrosomes are attenuated and ncMTOCs are active. At these stages, the polarization of the MT cytoskeleton is essential for the asymmetric localization of mRNAs and their encoded proteins, such as *gurken (grk)* and *oskar (osk)*, which are critical for the establishment of the embryonic axes^18^. Previous work showed that ncMTOCs are required for oocyte polarization, as Shot mutants show mislocalization of *grk*, *osk*^19^ and Staufen^14^, an RNA-binding protein that associates with *osk* and mediates its transport and anchoring at the posterior region^18^.

To determine whether Wdr62 is required for the asymmetric localization of oocyte determinants, Wdr62 was depleted during oogenesis using tissue-specific RNA interference (RNAi) (**Supplementary Fig. 2a-d**). RNAi expression was driven by the maternal α-tubulin-Gal4 driver, which becomes active during stage 2–3 egg chambers, after the initial four mitotic cycles and oocyte specification^48^. This strategy bypasses potential earlier requirements for Wdr62 during germline development.

In wild-type mid-stage oocytes, *grk* mRNA is transported by the minus-end-directed motor protein Dynein to the antero-dorsal side of the oocyte, near the nucleus, where it is subsequently translated^49^. Accordingly, in all control oocytes (mCherry RNAi), Grk protein localized normally to the antero-dorsal cortex, forming a crescent around the nucleus^50^ (**Fig. 2a**). As expected, based on previous reports^14^, Grk was mislocalized in oocytes depleted of Shot (**Fig. 2a, b**). Similarly, Wdr62-depleted oocytes showed mislocalization of Grk in 59% of oocytes analyzed compared to 0% in controls (**Fig. 2a, b**). Consistent with this, Grk levels at the antero-dorsal corner were significantly reduced in oocytes depleted of either Shot or Wdr62 (**Fig. 2a, c**). Consistent with defects in Grk localization, eggs laid by females depleted of Wdr62 in the germline frequently exhibited abnormalities in dorsal appendage morphology, including shortened, fused, or absent appendages (**Supplementary Fig. 3a, b**). As dorsal appendage formation depends on Gurken-mediated signaling to the surrounding follicle cells, these phenotypes are consistent with impaired Grk signaling in Wdr62-depleted oocytes.

**Figure 2.**
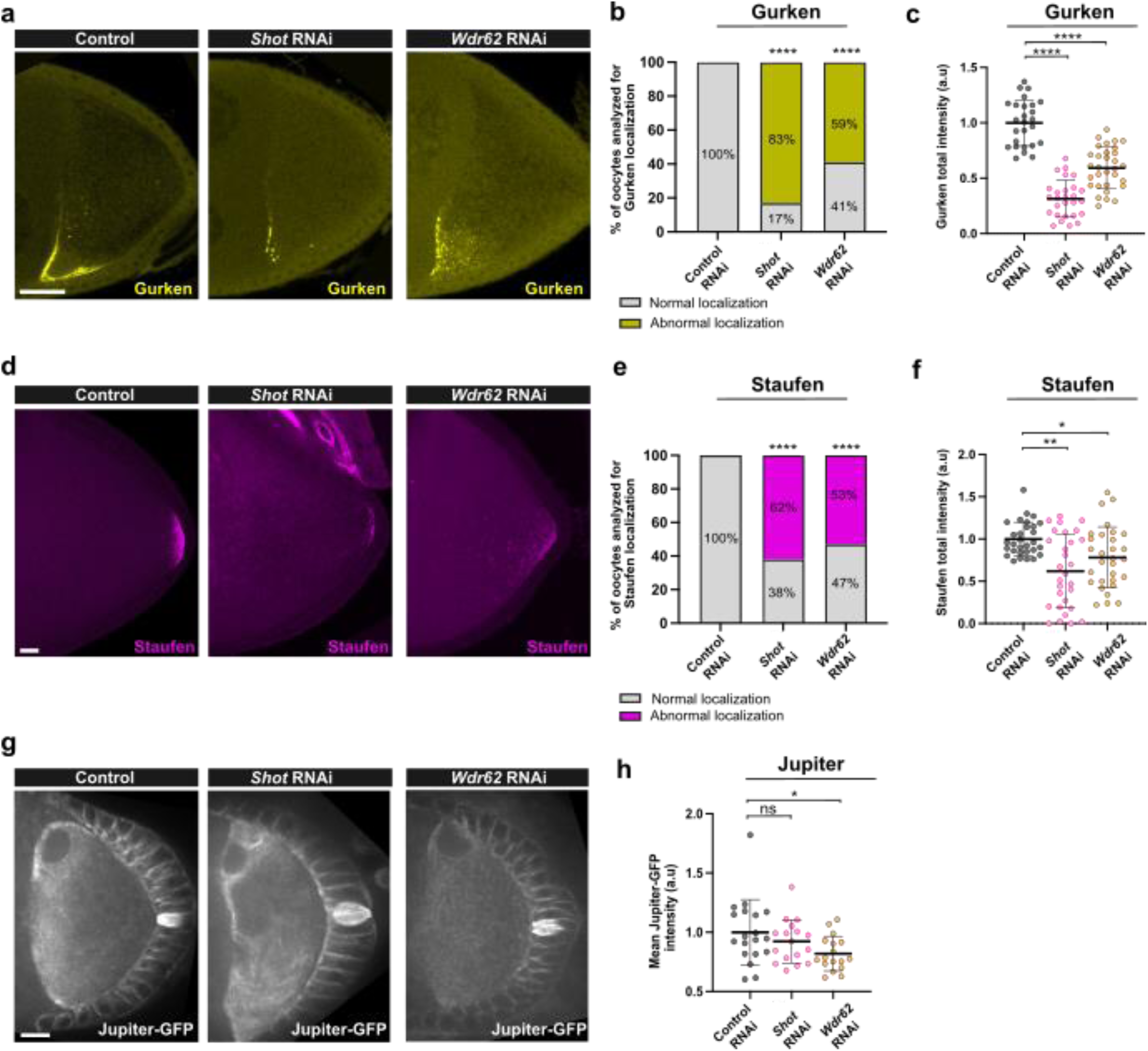
– Wdr62 is required for oocyte polarity and MT organization in the oocyte. **a**) Representative images of Grk localization at the antero-dorsal corner of mid-stage (stage 8) oocytes in control (mCherry RNAi), Shot RNAi, and Wdr62 RNAi conditions. **b**) Quantification of oocytes displaying normal or abnormal Grk localization for each condition. Values indicate the percentage of oocytes scored in each category. **c**) Quantification of total Grk fluorescence intensity at the antero-dorsal corner in control (mCherry RNAi, n = 26), Shot RNAi (n = 27) and Wdr62 RNAi (n = 31) conditions. **d**) Representative images of Staufen-RFP localization at the posterior pole of stage 9-10 oocytes in control (mCherry RNAi); Shot RNAi and Wdr62 RNAi conditions. **e**) Quantification of oocytes displaying normal or abnormal Staufen localization for each condition. Values indicate the percentage of oocytes scored for each category. **f**) Quantification of total Staufen fluorescence intensity at the posterior pole in control (mCherry RNAi, n = 30), Shot RNAi (n = 29) and Wdr62 RNAi (n = 30). **g**) Representative still images from live imaging of the MT network labeled with Jup-GFP in stage 7-8 oocytes, in control (mCherry RNAi), Shot RNAi and Wdr62 RNAi. **h**) Quantification of the mean fluorescence intensity (± SD) of Jupiter-GFP in control (mCherry-RNAi, n = 19), Shot RNAi (n = 17) and Wdr62 RNAi (n = 17). Statistical significance was assessed using the Fisher’s exact test (b,e) and the Kruskal–Wallis test with multiple comparisons (c,f,h) (*p < 0.05; **p < 0.01; ****p < 0.0001). Scale bars, 10 μm.

We next examined whether posterior determinants are affected upon Wdr62 loss. In control oocytes, Staufen localizes to a crescent at the posterior of the oocyte (**Fig. 2d**). As previously observed in *Shot* mutants^14^, Staufen failed to properly localize to the posterior region in Shot-depleted oocytes (62% vs. 100% in control; **Fig. 2d, e**). A similar defect was observed upon Wdr62 depletion, with 53% of oocytes exhibiting defective posterior Staufen localization (**Fig. 2d, e**). Consistent with this, in both Shot– and Wdr62-depleted oocytes, Staufen levels at the posterior cortex were reduced (**Fig. 2d, f**). Together, these results indicate that depletion of Wdr62 disrupts MT-dependent polarization of the oocyte.

To determine whether these defects reflect alterations in the underlying MT cytoskeleton, the overall organization of MTs was next examined in fixed oocytes by anti-a-tubulin staining and by still images of *in vivo* MT labeling using Jupiter-GFP (Jup-GFP), a MT-associated protein that binds along the MT lattice and serves as a reporter of MT abundance^51^. In control oocytes, MTs are enriched at the anterior cortex, forming a polarized network that progressively decreases in density toward the posterior^39^ (**Fig. 2g; Supplementary Fig**. **4**). Consistent with previous work^14^, Shot-depleted oocytes showed a redistribution of MTs towards the middle region of the oocyte compared with controls (**Fig. 2g, h; Supplementary Fig**. **4**). However, Wdr62-depleted oocytes showed an overall reduction in MT density throughout the oocyte cytoplasm compared to controls. This was confirmed by quantification of Jup-GFP fluorescence intensity, which revealed a significant decrease in Jup-GFP signal in Wdr62-depleted oocytes compared to controls (**Fig. 2g, h**), suggesting that Wdr62 contributes to the assembly and/or maintenance of the oocyte MT cytoskeleton.

Altogether, these observations suggest that Wdr62 contributes both to the polarized organization of the oocyte MT cytoskeleton and to the overall integrity of the MT network during mid-oogenesis.

### Wdr62 regulates the assembly of cortical non-centrosomal microtubule-organizing centres

Given that Wdr62 colocalizes with Shot and Patronin, and that its depletion disrupts oocyte polarization and MT organization, as observed in Shot and Patronin mutants, we next examined whether Wdr62 plays a role in ncMTOC assembly. To address this, Wdr62 was depleted and the localization of Shot and Patronin was analyzed. As shown in Figure 1, in control stage 9 oocytes, Shot and Patronin localize to ncMTOCs at the antero-lateral cortex, from which MTs have been shown to grow^14^. Loss of Wdr62 resulted in increased Shot accumulation at the antero-lateral cortex and within the oocyte cytoplasm, where Shot formed discrete foci (**Fig. 3a-c arrowhead**). Quantification of Shot fluorescence intensity revealed an approximately twofold increase at both the cortex and within the cytoplasm compared with controls (**Fig. 3b, c**). Analysis of a transgene expressing endogenous Patronin-YFP^14^ showed that Patronin localized robustly to the oocyte cortex in control oocytes (**Fig. 3d, e**). Patronin localization at the antero-lateral cortex was severely impaired upon Wdr62 depletion, with 100% of oocytes showing loss of cortical Patronin (**Fig. 3d, e**). Notably, this loss was more pronounced than in Shot-depleted oocytes (**Fig. 3d, e**), suggesting that Wdr62 may play a more prominent role in regulating Patronin localization.

**Figure 3.**
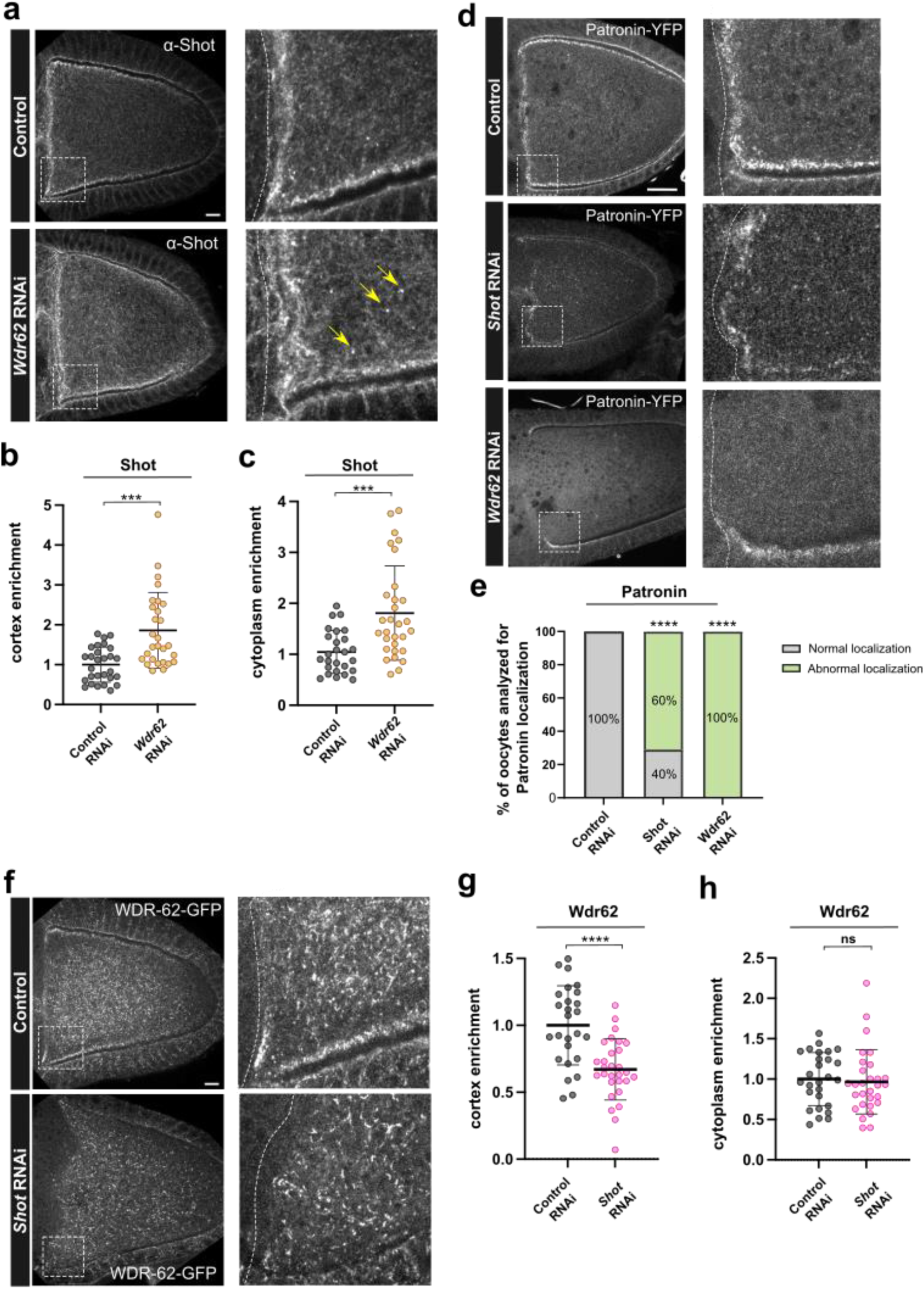
– Wdr62 is required for proper recruitment of Shot and Patronin at the oocyte cortex. **a**) Representative images of Shot localization in stage 9-10 oocytes in control (mCherry RNAi) and Wdr62 RNAi. Insets correspond to magnified views of boxed regions. Insets show higher-magnification views of the boxed regions. Note accumulation of Shot foci in the oocyte cytoplasm of Wdr62-depleted oocytes. **b**) Quantification of Shot mean fluorescence intensity at the antero-lateral cortex of control (mCherry RNAi, n = 28) and Wdr62 RNAi (n = 28) oocytes. **c)** Quantification of Shot mean fluorescence intensity within the cytoplasm of control (mCherry RNAi, n = 26) and Wdr62 RNAi (n = 30) oocytes. **d**) Representative images of Patronin::YFP localization in stage 9-10 oocytes in control (mCherry RNAi, n = 28), Shot RNAi (n = 25) and Wdr62 RNAi (n = 28) conditions. Insets correspond to magnified views of boxed regions. **e**) Quantification of oocytes displaying normal or abnormal Patronin::YFP localization in the different RNAi conditions. N = 28 for control (mCherry RNAi), n = 25 for Shot RNAi, and n = 28 for Wdr62 RNAi. Values indicate the percentage of oocytes scored for each category. **f**) Representative images of Wdr62::GFP localization in stage 9-10 oocytes in control (mCherry RNAi) and Shot RNAi conditions. Insets correspond to magnified views of boxed regions. **g**) Quantification of Wdr62::GFP mean cortical fluorescence intensity at the antero-lateral cortex in control (mCherry RNAi, n = 26) and Shot RNAi (n = 29) oocytes. **h**) Quantification of mean cytoplasmic Wdr62::GFP fluorescence in oocytes from control (mCherry RNAi) and Shot RNAi oocytes (mCherry RNAi, n = 27 and Shot RNAi, n = 29). Each dot represents one oocyte. Statistical significance was determined using the unpaired Mann–Whitney test or Fisher’s exact test, as appropriate (**p < 0.01; ***p < 0.001; ****p < 0.0001). Scale bars, 10 μm.

Shot has previously been shown to localize to the oocyte via its actin-binding domain and to recruit Patronin to cortical ncMTOCs^14^. We therefore asked whether Shot is also required for cortical localization of Wdr62, which could account for the loss of Patronin upon Wdr62 depletion. To address this, Shot was depleted by RNAi (**Supplementary Fig. 2e-g**), and the localization of Wdr62::GFP was analyzed. Depletion of Shot led to a significant reduction in cortical Wdr62 levels (**Fig. 3 f, g**), whereas Wdr62 levels within the oocyte cytoplasm were not affected (**Fig. 3f, h**). These observations suggest that Shot function is required for the cortical localization of Wdr62, but not for its cytoplasmic localization.

Together, these results show that Wdr62 depletion impairs the cortical localization of Patronin and altered accumulation and distribution of Shot at both cortical and cytoplasmic sites. Conversely, Shot depletion reduces cortical Wdr62 levels without affecting its cytoplasmic distribution. These observations demonstrate that Wdr62 and Shot are interdependent for proper cortical localization and that Wdr62 is required for proper assembly and composition of cortical ncMTOCs.

### Wdr62 is required for proper microtubule growth dynamics

To gain further insight into how Wdr62 regulates MT organization, MT dynamics were analyzed in control and Wdr62-depleted oocytes using EB1-GFP, which labels growing MT plus ends as fluorescent “comets” that track polymerizing MTs^52,53^. Shot-depleted oocytes were analyzed in parallel, as Shot is a ncMTOC component required for MT anchoring at the antero-lateral cortex^14^. Previous studies have shown that the dynamic behavior of MTs differs along the anterior–posterior axis^39,54^ (Cabrita *et al*., 2026 *in press*), which is consistent with the asymmetric distribution of cortical ncMTOCs. Because ncMTOCs are enriched at the antero-lateral cortex and excluded from the posterior, we reasoned that Wdr62-dependent defects might exhibit spatial specificity. Therefore, the oocyte was divided into three regions of interest: anterior, middle and posterior (**Fig. 4g**). EB1 comet number, lifetime (used as a measure of MT growth persistence), length, velocity and orientation were quantified within each region as previously done (Cabrita *et al*., 2026 *in press*).

**Figure 4.**
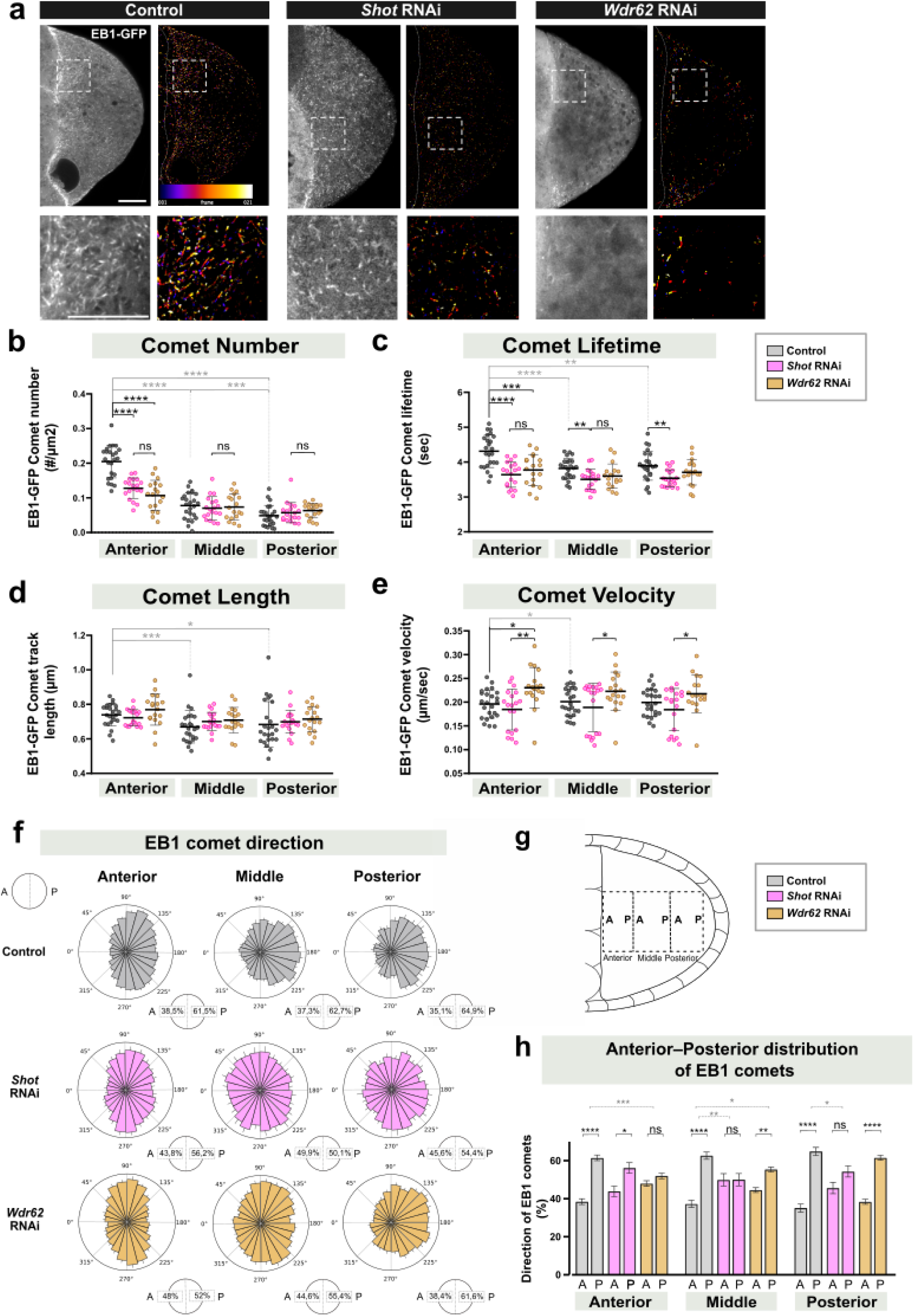
– Wdr62 depletion disrupts MT dynamics at the oocyte anterior region. **a**) Representative still images from live-imaging of EB1-GFP in control (mCherry RNAi), Shot RNAi and Wdr62 RNAi stage 8 oocytes. Left panel: maximum intensity projection of EB1-GFP signal at two consecutive time points. Right panel: EB1-GFP comet trajectories extracted from time-lapse movies using the Time-Lapse C plugin in FIJI to visualize growing MT plus ends. Temporal color coding (Temporal Color Code plugin) represents a time projection over 20 frames (0.5 s between frames). Insets show higher-magnification views of the corresponding boxed regions. **b-e)** Quantification of EB1-GFP comet dynamics obtained from time-lapse movies. For each quantified parameter, data represent mean ± SD for each region (anterior, middle and posterior) of stage 8 oocytes from control (mCherry RNAi, n = 24), Shot RNAi (n = 19) and Wdr62 RNAi (n = 18). Total comets analyzed per oocyte: 1431-2005. **b**) Scatter dot plot showing the mean number of EB1-GFP comets. **c**) Scatter dot plot showing the mean EB1-GFP comet lifetime. **d)** Scatter dot plot showing the mean EB1-GFP comet track length. **e**) Scatter dot plot showing the mean EB1-GFP comet velocity. For each parameter analyzed, one dot represents one oocyte for the corresponding region. **f**) Rose plots showing the orientation of EB1-GFP tracks within each region in control (mCherry RNAi, n = 24), Shot RNAi (n = 19) and Wdr62 RNAi (n = 18) oocytes. The mean percentage of tracks per oocyte oriented toward the anterior (A) or posterior (P) is indicated for each condition and region. **g**) Schematic representation of the oocyte regions used to analyze EB1-GFP comets dynamics in the oocyte for the different conditions. The oocyte was divided along the anterior-posterior axis into three regions: anterior, middle and posterior. For EB1 comet orientation analysis (h), within each region, EB1 comet tracks were classified based on their orientation toward the anterior (A) or the posterior (P). (**h**) Bar graph showing the mean percentage of EB1-GFP ± SEM comet track angles oriented toward the anterior and posterior sides of the oocyte. Percentages were calculated per oocyte and represent the relative distribution of comet orientation within each region. Statistical significance was assessed using one-way parametric ANOVA with Tukey’s multiple comparisons test or the Friedman test followed by Wilcoxon matched-pairs tests, as appropriate (see materials and methods) (\**p* < 0.05; **p < 0.01; ***p < 0.001; \*\*\*\**p* < 0.0001). Scale bars, 10 μm.

As previously reported by us and others (Cabrita *et al*., 2026 *in press*), control oocytes exhibit an enrichment of EB1 comets at the anterior region, with 0.205 ± 0.05 comets/μm^2^, followed by 0.078 ± 0.04 and 0.048 ± 0.03 comets/μm^2^ at the middle and posterior regions, respectively (**Fig. 4a, b**; **Table 1; Video 1**). In agreement with previous work^39^, we also observed a modest reduction in comet lifetime along the anterior-posterior axis, indicating reduced MT persistence toward the posterior (**Fig. 4c**). A similar decreasing trend was observed for comet length (**Fig. 4d**). Together, these measurements capture the progressive reduction in MT growth persistence and density toward the posterior and resolve region-specific differences in MT dynamics.

Consistent with a role for Shot in MT anchoring^14^, Shot depletion resulted in a significant reduction in EB1 comet density at the anterior region with 0.128 ± 0.03 comets/μm² (**Fig. 4a, b**; **Table 1; Video 2**). Similarly, Wdr62 depletion significantly reduced EB1 comet density at the anterior region with 0.107 ± 0.04 comets/μm², while comet density in the middle and posterior regions remained largely unchanged (**Fig. 4a, b**; **Table 1; Video 3**). These results suggest that both Shot and Wdr62 contribute to MT growth at the anterior cortex.

Shot and Wdr62-depleted oocytes exhibited reduced EB1 comet lifetimes at the anterior region, with an average of 3.64 s and 3.77 s, respectively, compared to 4.31 s in control oocytes (**Fig. 4c**; **Table 1**). A similar trend was observed in the middle and posterior regions, although only Shot depletion reached statistical significance (**Fig. 4c**). These results suggest that Wdr62 contributes to MT persistence at the anterior region, whereas Shot contributes to MT persistence throughout the oocyte.

EB1 comet length was not significantly affected by depletion of either Shot or Wdr62 in any region of the oocyte (**Fig. 4d**), indicating that the overall distance covered by growing MT plus ends is preserved. However, analysis of comet velocity revealed a significant increase in MT growth rate for Wdr62-depleted oocytes at the anterior region, with 0.230 ± 0.04 μm/s compared to 0.196 ± 0.03 μm/s in control oocytes (**Fig. 4e**; **Table 1**). Accordingly, comet velocity was significantly higher in Wdr62-depleted oocytes compared to Shot depletion in the middle and posterior regions. Together with the reduced comet number and lifetime, these results suggest that loss of Wdr62 results in faster but less persistent MT growth, consistent with a role in stabilizing MT dynamics.

Given that MT polarity underlies directed transport in the oocyte, we next examined EB1 comet directionality as a readout of MT network organization along the anterior–posterior axis. As previously reported, control oocytes exhibited a bias in comet orientation toward the posterior, consistent with the polarized organization of the MT network^39^ (**Fig. 4f, h**). Upon Wdr62 depletion, this directional bias was specifically disrupted at the anterior region (**Fig. 4f, h**). This is consistent with the alterations observed in the other MT dynamic parameters, suggesting that less persistent MT growth may reduce the ability of MTs to maintain directional trajectories at the anterior cortex. Interestingly, Shot depletion resulted in a loss of directional bias in both the middle and posterior regions of the oocyte (**Fig. 4f, h**), suggesting that Shot is required for maintaining MT growth orientation more broadly across the oocyte.

Together, these results show that Wdr62 regulates MT growth dynamics at the anterior cortex, contributing to MT stability and directional organization, whereas Shot acts more broadly across the oocyte. These findings support a role for Wdr62 in maintaining MT network polarity during oogenesis.

### Wdr62 functions with Patronin and cooperates with Katanin to regulate acentrosomal microtubule organization

Given that Wdr62 depletion severely disrupts Patronin enrichment at cortical ncMTOCs and alters MT growth dynamics, we next asked whether Wdr62 functionally interacts with known regulators of acentrosomal MT organization. Patronin is a well-established regulator of MT minus-end stability^20–22^, and proteomic analyses of *Drosophila* ovary extracts have shown that Patronin co-immunoprecipitates with the conserved MT-severing enzyme Kat80^14^. Because Katanin has been implicated in acentrosomal MT organization in multiple cell types^25^, we tested genetic interactions with Patronin and Katanin to determine whether they cooperate with Wdr62 in regulating MT dynamics in the oocyte.

Because no germline RNAi line is available to deplete Patronin, and strong *patronin* null mutants arrest oogenesis at early stages^14^, genetic interactions were tested using a heterozygous *patronin* mutant background carrying the hypomorphic allele, *patronin*^05252^, which strongly reduces Patronin levels^55^. The impact of combining Wdr62 depletion with reduced Patronin dosage was assessed by analyzing EB1 comet dynamics.

Previous work showed that *patronin*^05252^ homozygous oocytes contain 90% fewer cortical EB1-GFP foci^14^, whereas heterozygous mutants exhibit ∼50% fewer EB1-GFP comets in MT polymerization–depolymerization assays (Cabrita *et al*., 2026 *in press*). Consistent with this, heterozygous *patronin*^05252^ mutants exhibited a mild reduction in EB1 comet number, most apparent at the anterior region (0.166 ± 0.05 comets/μm^2^ compared to 0.194 ± 0.06 comets/μm^2^ in control) **(Supplementary Fig. 5a; Fig. 5b**; **Table 2; Video 4)**. The other parameters, were largely unchanged compared to controls, suggesting that reducing Patronin levels has a modest effect on MT growth dynamics.

**Figure 5.**
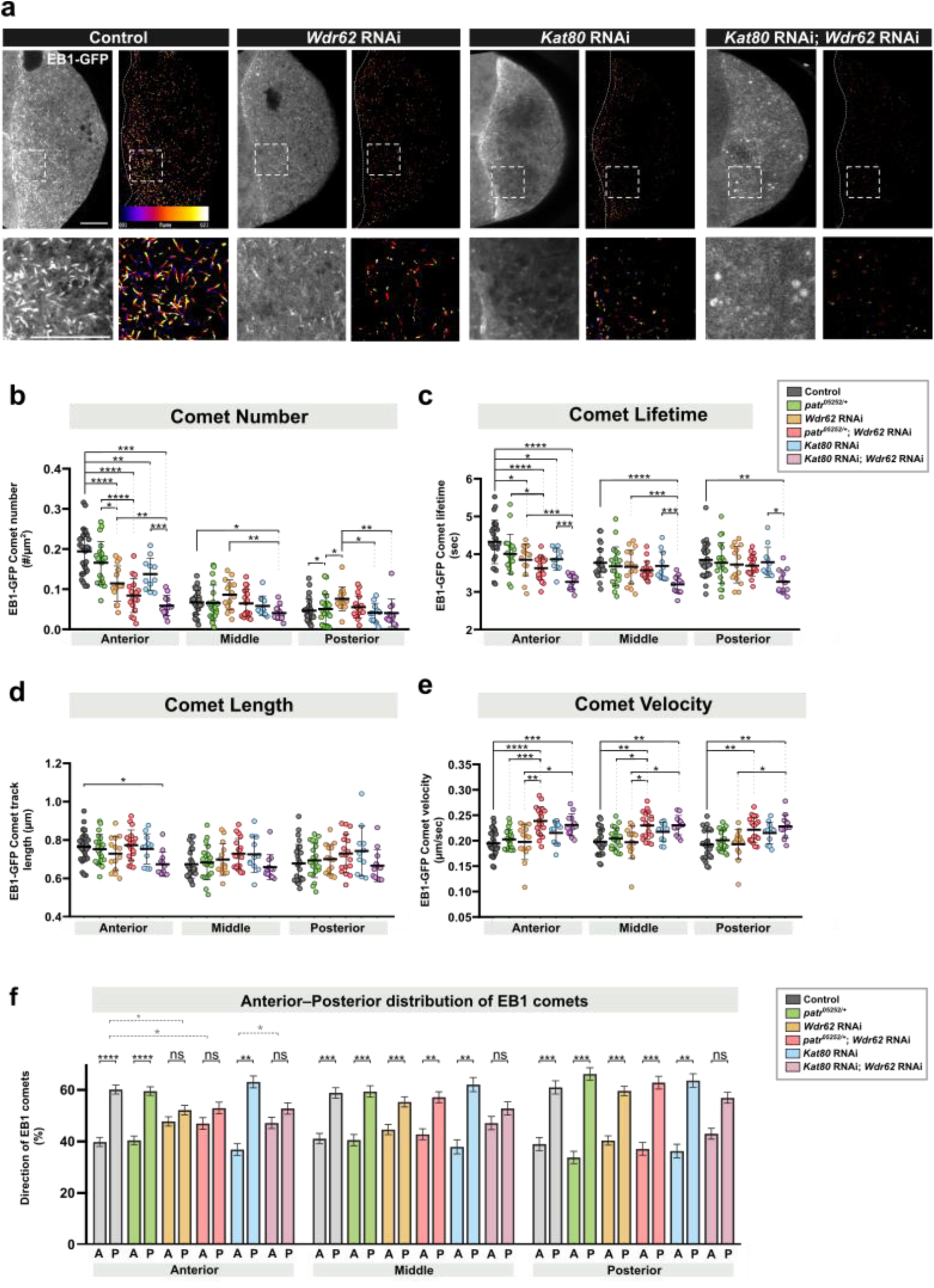
– MT dynamics upon combined perturbation of Wdr62 with Patronin or Katanin. **a**) Representative still images from live-imaging of EB1-GFP in stage 8 oocytes from control (mCherry RNAi), Wdr62 RNAi, Kat80 RNAi and combined Kat80 and Wdr62 RNAi. Left panel: maximum intensity projection of EB1-GFP signal at two consecutive time points. Right panel: EB1-GFP comet trajectories extracted from time-lapse movies using the Time-Lapse C plugin in FIJI to visualize growing MT plus ends. Temporal color coding (Temporal Color Code plugin) represents a time projection over 20 frames (0.5 s between frames). Insets show higher-magnification views of the corresponding boxed regions. **b-e)** Quantification of EB1-GFP comet dynamics obtained from time-lapse movies. For each quantified parameter, data represent mean ± SD for each region (anterior, middle and posterior) of stage 8 oocytes from control (mCherry RNAi, n = 27), *patr*^05252/+^ (n = 20), Wdr62 RNAi (n = 14), *patr*^5252/+^, Wdr62 RNAi (n = 18), Kat80 RNAi (n = 11) and combined Kat80 and Wdr62 RNAi (n = 12). Total comets analyzed per oocyte: 1025-1807. **b**) Scatter dot plot showing the mean number of EB1-GFP comets. **c**) Scatter dot plot showing the mean EB1-GFP comet lifetime. **d**) Scatter dot plot showing the mean EB1-GFP comet track length. **e**) Scatter dot plot showing the mean EB1-GFP comet velocity. For each parameter analyzed, one dot represents one oocyte for the corresponding region. **f)** Bar graph showing the mean percentage (± SEM) of EB1.GFO comet track angles oriented toward the anterior and posterior sides of the oocyte. Percentages were calculated per oocyte and represent the relative distribution of comet orientation within each region. Statistical significance was assessed using one-way ANOVA with Tukey’s multiple-comparisons test, Games–Howell test or Kruskal–Wallis test with multiple comparisons, as appropriate (*p < 0.05; **p < 0.01; ***p < 0.001; ****p < 0.0001). Scale bars, 10 μm.

Combining heterozygous *patronin*^05252^ mutants with Wdr62 depletion did not significantly enhance the reduction in EB1 comet density relative to Wdr62 depletion alone (0.084 ± 0.04 comets/μm^2^ vs 0.114 ± 0.04 comets/μm^2^ in Wdr62 RNAi) **(Supplementary Fig. 5a; Fig. 5b**; **Table 2; Video 5)**. A similar trend was observed for EB1 comet lifetime **(Fig. 5c**; **Table 2)**. Consistent with our previous experiments, EB1 comet velocity in Wdr62-depleted oocytes in the anterior region showed a modest increase, although this difference was not statistically significant different compared to controls. Comet velocity was further increased when both *patronin*^05252^ mutants were combined with Wdr62 RNAi (**Fig. 5e**; **Table 2**).

These results show that partial reduction of Patronin does not exacerbate the Wdr62 depletion phenotype. Together with the loss of Patronin in Wdr62-depleted oocytes, these findings suggest that Wdr62 and Patronin function within the same pathway to regulate MT organization in the oocyte.

We next examined the contribution of Kat80 to MT dynamics. Similarly to Wdr62 depletion, Kat80-depleted oocytes exhibited a significant reduction in EB1 comet number at the anterior region (0.137 ± 0.04 comets/μm^2^ compared to 0.194 ± 0.06 comets/μm^2^ in control and 0.114 ± 0.04 comets/μm^2^ in Wdr62-RNAi; **Fig. 5a, b**; **Table 2; Video 6**). This was accompanied by a decrease in comet lifetime at the anterior region, while comet length, velocity and orientation were largely unchanged (**Fig. 5c-f; Supplementary Fig. 5b**). These results indicate that Kat80 contributes to MT dynamics at the anterior region of the oocyte by affecting the number and persistence of MT growth events.

Strikingly, simultaneous depletion of Wdr62 and Kat80 resulted in a strong enhancement of the MT phenotype, with significant alterations across all MT dynamic parameters analyzed (**Fig. 5**; **Table 2; Video 7**), indicating a strong genetic interaction between the two proteins. EB1 comet density at the anterior region was significantly reduced when compared with either Wdr62 or Kat80 single depletions (0.059 ± 0.03 comets/μm^2^ compared to 0.114 ± 0.04 comets/μm^2^ in Wdr62-RNAi and 0.137 ± 0.04 comets/μm^2^ in Kat80-RNAi; **Fig. 5a, b**; **Table 2**). Comet lifetime was also further reduced in both anterior and middle regions, indicating a pronounced decrease in MT growth persistence (**Fig. 5c**; **Table 2**). In addition, MT growth directionality was strongly disrupted, with loss of posterior bias in both anterior and middle regions (**Fig. 5f; Supplementary Fig.5b**), whereas this bias was only affected at the anterior region in Wdr62 depletion and remained largely unchanged in Kat80-depleted oocytes (**Fig. 5f; Supplementary Fig.5b**). Comet velocity in the double perturbation was significantly increased compared to control and Wdr62-depleted oocytes, while not differing significantly from Kat80 depletion alone (**Fig. 5e**). Importantly, the combined depletion of Wdr62 and Kat80 resulted in a markedly stronger reduction in EB1 comet number, lifetime and length than observed in either single condition (**Fig. 5 b-d**). These results indicate that MT growth is severely compromised in oocytes lacking both Wdr62 and Kat80. Moreover, these findings reveal a synergistic interaction between Wdr62 and Kat80 in regulating MT dynamics.

Together these findings suggest that Wdr62 and Patronin function in the same pathway, and that Wdr62 functionally cooperates with Kat80 to regulate MT dynamics and organization during mid-oogenesis.

### Wdr62 regulates acentrosomal meiotic spindle assembly and fertility through cooperation with Patronin

Given that Wdr62 regulates acentrosomal MT organization during mid-oogenesis, we next asked whether Wdr62 also contributes to MT organization in another acentrosomal context during oogenesis, such as, the assembly of the meiotic spindle. In *Drosophila* oocytes, centrosomes are progressively eliminated, and the meiotic spindle is assembled through acentrosomal mechanisms^38^. We therefore examined whether Wdr62 localizes to and functions within the meiotic spindle. Analysis of endogenously tagged Wdr62::GFP revealed that Wdr62 localizes to metaphase I spindles, where it is distributed along spindle MTs (**Fig. 6a**).

**Figure 6.**
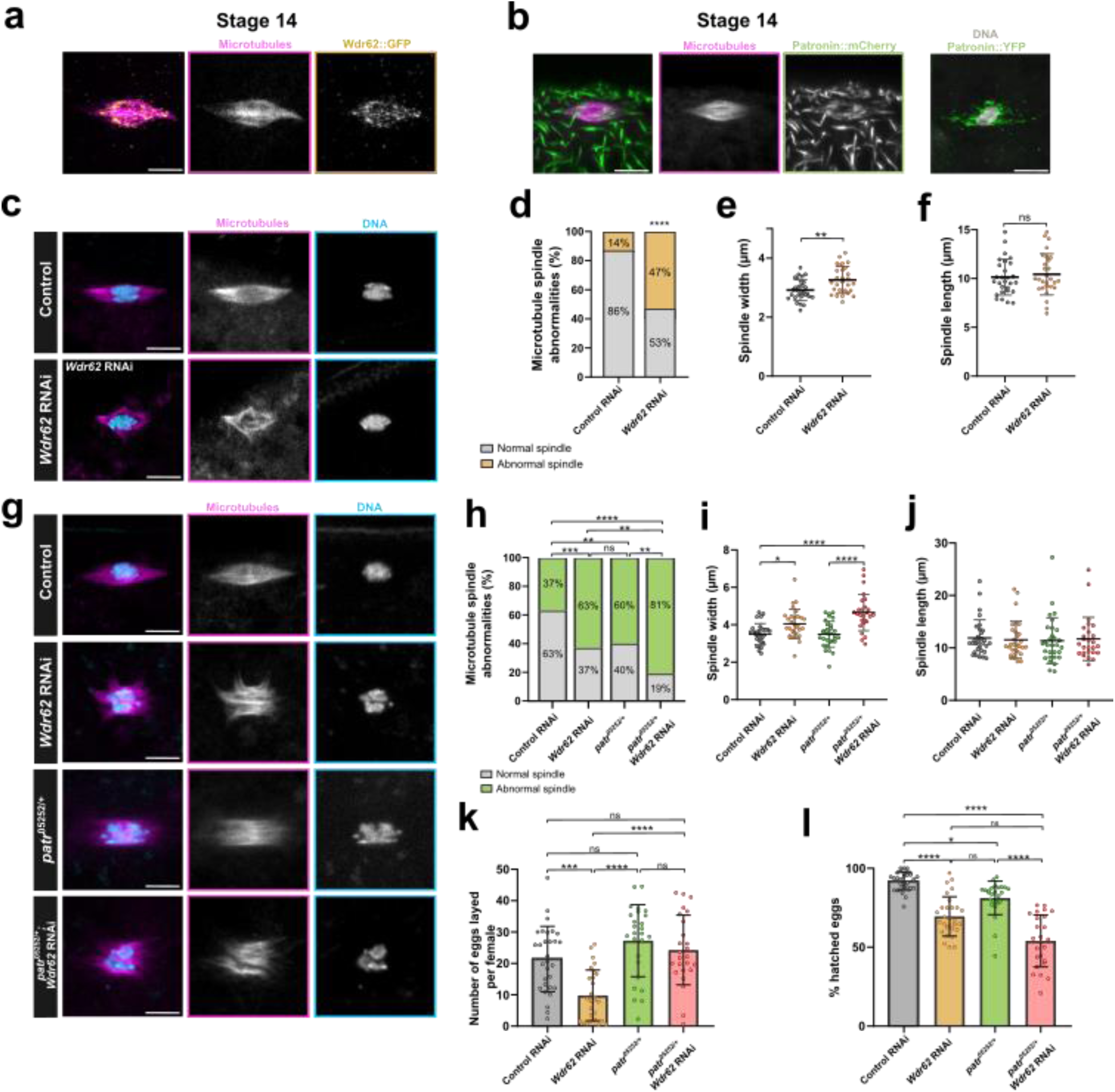
– Wdr62 and Patronin are required for oocyte spindle assembly and fertility. (**a**) Representative images of Wdr62::GFP (yellow) localization along MTs (Jup-GFP; magenta) of the metaphase I oocyte spindle in stage 14 oocytes. **b**) Representative images Patronin::mCherry or Patronin::YFP (green) localization along MTs (Jup-GFP; magenta) of the metaphase I oocyte spindle in stage 14 oocytes. **c)** Representative images of metaphase I spindles in control (mCherry RNAi) and Wdr62 RNAi conditions. MTs (Jup-GFP; magenta) and DNA (Cyan). **d**) Quantification of MT spindle morphology in control (mCherry RNAi, n = 39) and Wdr62 RNAi (n = 28) showing the proportion of oocytes displaying normal and abnormal spindle morphology. Values indicate the percentage of spindles scored for each category. **e)** Quantification of MT spindle width in control (mCherry RNAi, n = 39) and Wdr62 RNAi (n = 28). **f**) Quantification of MT spindle length in control (mCherry RNAi, n = 39) and Wdr62 RNAi (n = 28). **g)** Representative images of metaphase I spindles in control (mCherry RNAi), Wdr62 RNAi, *patr*^05252/+^ and *patr*^05252/+^; Wdr62 RNAi oocytes. MTs (Jup-GFP; magenta) and DNA (Cyan). **h**) Quantification of MT spindle morphology in control (mCherry RNAi, n = 30), Wdr62 RNAi (n = 30), *patr*^05252/+^ (n = 30) and *patr*^05252/+^; Wdr62 RNAi (n = 26) oocytes showing the proportion of oocytes displaying normal and abnormal spindle morphology for each condition. Values indicate the percentage of spindles scored for each category. **i**) Quantification of MT spindle width in control (mCherry RNAi, n = 30), Wdr62 RNAi (n = 30), *patr*^05252/+^ (n = 30) and *patr*^05252/+^; Wdr62 RNAi (n = 26) oocytes. **j**) Quantification of MT spindle length in control (mCherry RNAi, n = 30), Wdr62 RNAi (n = 30), *patr*^05252/+^ (n = 30) and *patr*^05252/+^; Wdr62 RNAi (n = 26) oocytes. (**k**) Quantification of the number of eggs laid by control (mCherry RNAi, n = 30), Wdr62 RNAi (n = 31), *patr*^05252/+^ (n = 26) and *patr*^05252/+^;Wdr62 RNAi (n = 25) females. (**l**) Quantification of the percentage of hatched eggs in control (mCherry RNAi, n = 30), Wdr62 RNAi (n = 31), *patr*^05252/+^ (n = 26) and *patr*^05252/+^;Wdr62 RNAi (n = 25) conditions. Statistical significance was tested by Chi-square test, t-test and Kruskal–Wallis multiple comparisons test, as appropriate (*p < 0.05; **p < 0.001; ***p < 0.0001). Scale bars, 5 μm.

To assess whether Wdr62 is required for spindle organization, meiotic spindle morphology was examined in Wdr62-depleted oocytes expressing Jup-GFP, which labels spindle MT^51^. In control oocytes, the metaphase I spindle adopts a bipolar fusiform structure with focused poles and chromosomes aligned at the equator (**Fig. 6c**). In contrast, Wdr62-depleted oocytes frequently exhibited abnormal spindle morphologies, including unfocused poles and frayed MTs (**Fig. 6c, g**). Quantification revealed that 47% of Wdr62-depleted oocytes exhibited abnormal spindle morphology, compared with 14% of control oocytes (**Fig. 6d**). Measurement of spindle dimensions further showed that Wdr62 depletion significantly increased spindle width (**Fig. 6e**), whereas spindle length remained unchanged (**Fig. 6f**). Together, these results demonstrate that Wdr62 is required for proper organization of the acentrosomal meiotic spindle in oocytes.

In addition to spindle defects, Wdr62 depletion also delayed nuclear envelope breakdown (NEBD) during stage 13. In control oocytes, NEBD was observed in 99% of late stage-13 oocytes, whereas only 76% of Wdr62-depleted oocytes had undergone NEBD at this stage (**Supplementary Fig. 6a, b**). This delay suggests that Wdr62 also contributes to the timely progression of meiotic maturation. Consistent with this, Wdr62-depleted ovaries contained fewer stage 14 eggs (**Supplementary Fig. 6c)**.

Having established a requirement for Wdr62 in meiotic spindle organization, we next examined whether Wdr62 and Patronin cooperate during this process. To assess Patronin localization during meiosis, stage-14 oocytes expressing endogenously tagged Patronin–YFP were analyzed. The YFP signal outlined a bipolar structure consistent with the metaphase I spindle morphology **(Fig. 6b)**. Co-expression of Patronin–mCherry with Jup–GFP further confirmed that Patronin decorates spindle MTs **(Fig. 6b)**. Functional interaction between Wdr62 and Patronin was then tested genetically. Spindle morphology was examined in Wdr62-depleted oocytes, *patronin*^05252^ heterozygous mutants, and in oocytes combining both perturbations.

As previously observed, Wdr62-depleted oocytes frequently displayed spindle abnormalities (63% abnormal spindles compared with 37% in controls; **Fig. 6g, h**). Notably, *patronin*^05252^ heterozygous mutants also exhibited spindle defects, with 60% abnormal spindles, indicating that reduced Patronin levels compromise meiotic spindle organization. Strikingly, combining Wdr62 depletion with *patronin*^05252^ heterozygous mutants further increased the frequency of abnormal spindles to 81%, significantly exceeding either perturbation alone (**Fig. 6g, h**). Consistent with this, analysis of spindle dimensions showed that Wdr62 depletion significantly increased spindle width, whereas *patronin*^05252^ heterozygous mutants alone did not significantly alter this parameter (**Fig. 6g, i**). Combined perturbation further increased spindle width relative to controls and Patronin heterozygous mutants, although the difference relative to Wdr62 depletion alone did not reach statistical significance (**Fig. 6g, i**). Spindle length remained unchanged in all conditions analyzed (**Fig. 6g, j**).

To assess the functional requirement of Wdr62 and Patronin during oogenesis and their impact on female fertility, egg production and hatching rates were quantified in females depleted of Wdr62, in *patronin*^05252^ heterozygous mutants, and in the combined genetic background. Females depleted of Wdr62 laid significantly fewer eggs than controls (**Fig. 6k**). In addition, egg viability was strongly compromised, as the percentage of eggs that hatched was significantly reduced compared with controls (**Fig. 6l**), indicating that Wdr62 is required for both egg production and embryo viability. Patronin heterozygous mutant females produced a similar number of eggs as controls (**Fig. 6k**), however the number of hatched eggs was significantly reduced in comparison with controls (**Fig. 6l**). Interestingly, females combining Wdr62 depletion with reduced Patronin dosage laid more eggs than Wdr62-depleted females alone **(Fig. 6k**). Although the basis of this effect remains unclear, one possibility is that partial reduction of Patronin levels alleviates the delay observed upon Wdr62 depletion in NEBD, thereby allowing more oocytes to progress to late stages and be laid as eggs. Despite this partial recovery in egg production, egg viability was not improved, as combining Wdr62 depletion with reduced Patronin dosage did not significantly further reduce hatching rates relative to the single perturbations, although a trend toward reduced hatching was observed (**Fig. 6l**).

Collectively, these results indicate that the conserved proteins Wdr62 and Patronin are required for meiotic spindle assembly and female fertility, and that their cooperative function during oogenesis ensures proper oocyte development and reproductive success.

## DISCUSSION

Our study identifies Wdr62 as a conserved regulator of acentrosomal MT organization across multiple stages of *Drosophila* oogenesis. We show that Wdr62 localizes to cortical ncMTOCs in mid-oogenesis and is required for the proper recruitment of the core ncMTOC components Shot and Patronin. Loss of Wdr62 disrupts oocyte polarization, alters MT dynamics during mid-oogenesis, and impairs meiotic spindle assembly, indicating that Wdr62 contributes broadly to the organization of acentrosomal MT networks. Consistent with these roles, depletion of Wdr62 or Patronin compromises fertility outcomes.

Throughout oogenesis, Wdr62 preferentially localizes to regions where MTs are generated through acentrosomal mechanisms. These include the fusome, an actin-rich structure essential for oocyte determination in the germarium that contains acetylated MTs^56^, the antero-lateral cortex of mid-stage oocytes, and the apical surface of follicular epithelial cells, to which Shot and Patronin also localize^14,42,47,56^, as well as the oocyte spindle, where we found that both Patronin and Wdr62 localize. In early stages of oogenesis, when centrosomes are still active as MTOCs, Wdr62 predominantly colocalizes with Shot and Patronin rather than centrosomes, supporting a conserved central role in acentrosomal MT organization.

Our observations are consistent with work in *C. elegans* intestinal epithelial cells, where WDR-62 localizes to apical ncMTOCs, interacts with the Patronin homologue PTRN-1, and is required for recruitment of both PTRN-1 and the Shot homologue VAB-10B^34^. In this context, WDR-62 acts upstream of these components. We found that depletion of Wdr62 in *Drosophila* oocytes led to a near-complete loss of Patronin but an increased accumulation of Shot. These results suggest that, although Wdr62 functions as a ncMTOC component in both systems, the mechanisms by which it regulates ncMTOC assembly are context dependent.

Similar variability has been reported for other ncMTOC components, supporting the view that ncMTOC composition and regulation are context dependent and can differ across cellular and developmental contexts^14,57,58^. These observations highlight the importance of investigating the function of ncMTOC components across diverse cell types to fully understand how acentrosomal MT networks are assembled and regulated.

Consistent with a role in ncMTOC function, loss of Wdr62 disrupts MT-dependent oocyte polarization, leading to mislocalization of the embryonic axis determinants Gurken and Staufen, as observed in Shot^19^ and Patronin^14^ mutants. Analysis of EB1 comets further supports the conclusion that MT dynamics are perturbed in Wdr62-depleted oocytes, with reduced EB1 comet number and lifetime, and altered orientation of EB1 comets at the anterior cortex. Consistent with this, WDR-62 depletion in *C. elegans* intestinal epithelial cells similarly reduces the number and length of growing MTs^34^, supporting a conserved role in promoting persistent MT growth.

Shot has been described as an anchor of ncMTOCs at the oocyte cortex by recruiting Patronin^14^. Our observations extend this model, as Shot alters MT dynamics throughout the oocyte, reducing MT growth persistence and perturbing MT orientation. This broader effect, together with the observed mesh-like distribution of Shot in the oocyte cytoplasm and the presence of a cytoplasmic actin mesh required for MT organization^44^, suggests that Shot contributes to MT stabilization beyond cortical ncMTOCs. Such a role is consistent with previous work in neurons showing that Shot can guide and stabilize MT growth along cortical actin networks, where Shot depletion similarly results in reduced EB1 comet lifetime and direction^59^. Together, these observations suggest that Shot may coordinate MT organization both at cortical ncMTOCs and within the oocyte cytoplasm.

Given that Wdr62 and Patronin co-localize at cortical ncMTOCs, and that Wdr62 depletion leads to loss of Patronin from the cortex, our data suggest that Wdr62 may regulate MT organization through Patronin-dependent mechanisms. Patronin is a conserved MT minus-end stabilizer that protects MTs from depolymerization and promotes the formation of stable acentrosomal MT arrays^20–22^. Consistent with this, partial reduction of Patronin dosage did not significantly enhance the Wdr62 depletion phenotype, suggesting that Wdr62 and Patronin likely function within the same pathway to regulate MT stability in the oocyte.

Given the large size of the oocyte and the density of its MT network, stabilization of MT minus ends alone is unlikely to fully account for the generation of this network. Canonical MT nucleation mediated by γ-tubulin is unlikely to contribute at cortical ncMTOCs, as γ-tubulin is not detected at these sites^14^, suggesting that additional mechanisms are required to sustain the generation of MTs. In other cellular contexts without centrosomes, such as neurons, MT amplification can be mediated by severing enzymes such as Katanin, which generate new MT seeds through severing of pre-existing MTs, thereby increasing MT number^25^. Kat80 has been shown to localize to the oocyte cortex, supporting a role at cortical ncMTOCs^14^. Consistent with this, we observed that Kat80-depleted oocytes show a significant reduction in EB1 comet number and lifetime at the anterior region. These observations show that Katanin contributes to MT organization in the oocyte, likely promoting the generation of MT growth events.

Notably, simultaneous depletion of Wdr62 and Kat80 resulted in a strong disruption of MT dynamics compared to either single depletion. These enhanced phenotypes suggest that Wdr62 and Kat80 functionally cooperate to regulate MT dynamics and organization in oocytes. Consistent with this, WDR62 and Katanin physically interact in human cells, where WDR62 recruits Katanin to spindle poles to regulate MT minus-end depolymerization, a process required for proper chromosome segregation^27,28^. Both *Drosophila* and human WDR62 preferentially bind curved MTs of cultured human cells or *in vitro* stabilized MTs, where WDR62 recruited Katanin for MT severing^27^. The interaction between WDR62 and subunit p80 of Katanin was shown to enhance the severing activity of Katanin *in vitro*, suggesting that WDR62 specifically targets Katanin to sever extended MT lattices^28^. Consistent with this previous work, the localization of Wdr62 in the oocyte cytoplasm also resembles what could be curved MTs (**Fig. 1g**, arrowhead), raising the possibility of a similar function for Wdr62 in the oocyte cytoplasm. Indeed, Kat80 has been shown to co-immunoprecipitate with Patronin from *Drosophila* ovary extracts^14^. Collectively, our data support a mechanism in which Wdr62, together with Patronin, stabilizes MT minus ends and may promote the recruitment of Katanin for MT severing. In this context, severing would generate new MT fragments whose exposed minus ends are subsequently stabilized by Patronin in a Wdr62-regulated manner, thereby coupling MT severing with minus-end protection and enabling sustained MT growth. Together, these observations support a model in which acentrosomal MT network assembly relies on the coordination of stabilization and amplification, with Wdr62 potentially acting as a coupling factor between these processes (**Fig. 7**). Interestingly, a recent preprint in *C. elegans* neurons suggests that PTRN-1 and the WD40-repeat protein, WDR-47, which are known to interact, localize to sites of MT severing mediated by Katanin. These data further suggest that both proteins may remain associated with newly generated MT minus ends following severing events, thereby stabilizing these minus ends. Although Wdr62 is distinct from WDR47, it also contains WD40-repeat domains, raising the possibility that Wdr62 may function in an analogous manner in the *Drosophila* oocyte.

**Figure 7.**
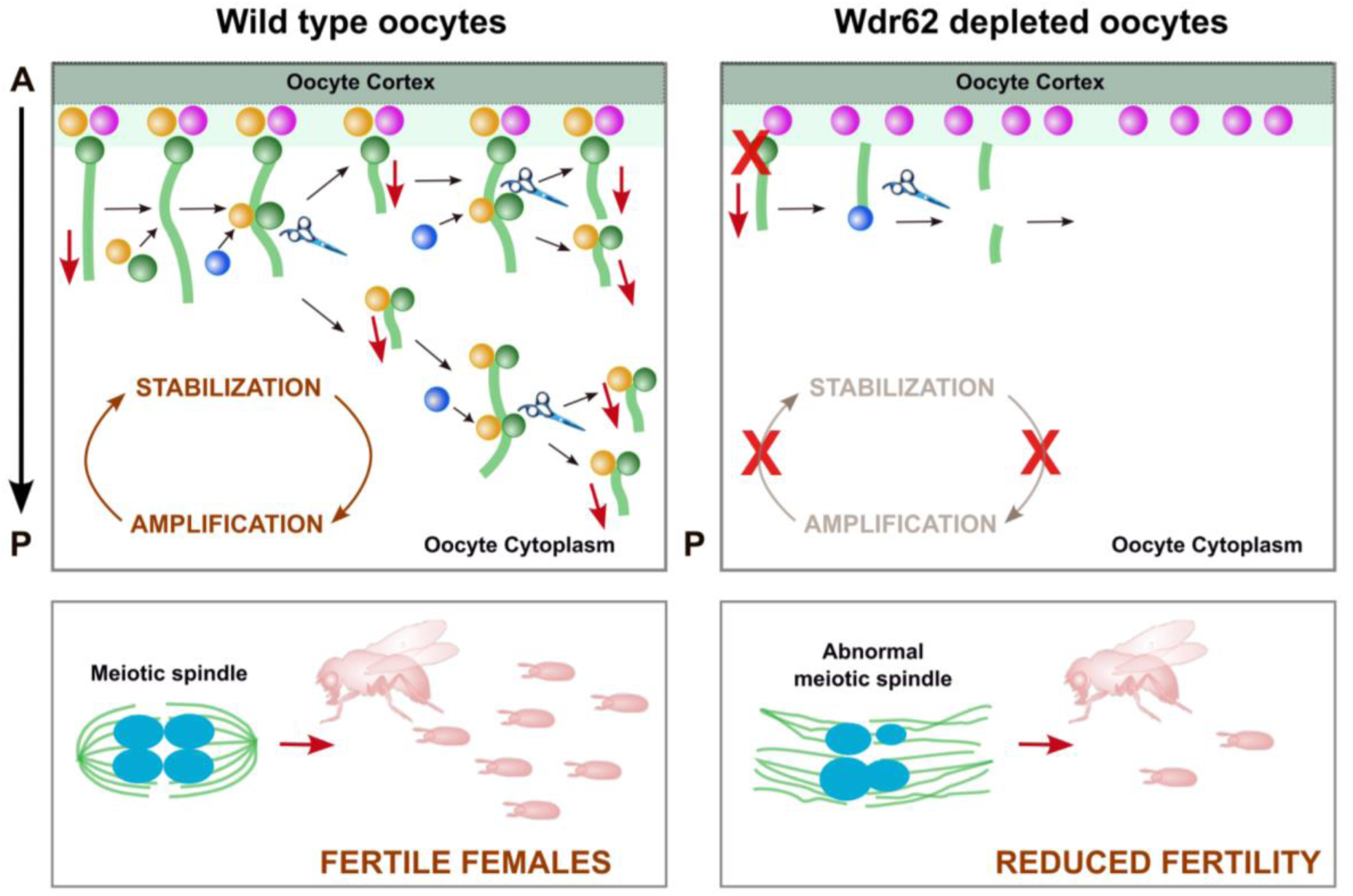
– Model of the stabilization–amplification Cycle for acentrosomal microtubule organization in the *Drosophila* oocyte. The assembly of the MT network is proposed to rely on the coordinated balance between stabilization and amplification, with Wdr62 potentially acting as a coupling factor between these processes. By working together with Patronin, Wdr62 promotes stabilization of MT growth events and may contribute to Katanin-mediated MT severing. Katanin-mediated severing generates new MT fragments, whose minus ends are stabilized by Wdr62 and Patronin, and from which MT outgrowth proceeds. This Stabilization-Amplification Cycle allows the assembly of the dense MT network in oocytes, which is critical for oocyte polarization. Loss of Wdr62 disrupts this cycle, which impacts oocyte polarization, alters MT dynamics, leads to abnormal spindle morphology, and ultimately compromises fertility.

Previous work showed that WDR62 is required for meiotic initiation in mouse oocytes, where its depletion leads to defects in meiotic progression, abnormal spindle morphology and increased incidence of aneuploidy^60,61^, the major cause of female infertility^62^. While these defects have been linked in part to altered JNK signaling, the cellular mechanisms underlying WDR62 function in oocyte meiosis remained incompletely understood^60^. Here, we show for the first time that Wdr62 localizes along MTs of oocyte spindles. Consistent with the mouse phenotype^60^, Wdr62 depletion in *Drosophila* oocytes results in abnormal spindle morphology, impaired meiotic progression and reduced fertility, supporting a conserved role for WDR62 in oocyte spindle assembly.

Building on this, we identify for the first time, the conserved MT minus-end stabilizer Patronin as an important regulator of oocyte spindle assembly^20,22,63^. Patronin, a core ncMTOC component in multiple cell types^14–16,64^, localizes to the oocyte spindle and is required for its proper organization and for fertility outcomes. Our data further shows that Patronin and Wdr62 functionally interact during this process. One possibility is that they stabilize distinct or overlapping populations of spindle MTs, which may explain the enhanced spindle defects observed upon their combined perturbation.

Katanin-mediated MT severing has been implicated in *C.elegans* oocyte spindle assembly, where it functions to amplify the MT spindle mass^65,66^. In mice, Katanin is also required for oocyte spindle assembly and fertility^60^, supporting a conserved role for MT severing in oocyte meiosis. Whether WDR62 and the MT minus-end stabilizer Patronin also cooperate with Katanin during meiotic spindle assembly remains unknown. Our findings provide a framework for future studies to test whether Katanin-mediated severing contributes to oocyte spindle assembly in *Drosophila* oocytes and, if so, to dissect how MT stabilization and amplification are coordinated to support this process.

Importantly, the role of Wdr62 in regulating the MT cytoskeleton during oogenesis has functional implications at the organismal level. Wdr62-depleted females exhibit reduced egg laying and decreased hatch rates, indicating compromised fertility. This aligns with observations in humans, where mutations in WDR62 are associated with premature ovarian insufficiency (POI)^67^, suggesting that disruption of WDR62 function in acentrosomal MT organization may impair oocyte development, thereby reducing oocyte competence and fertility.

Finally, given that WDR62 has been implicated in diverse developmental processes, including neuronal differentiation, brain development, microcephaly^29,30,68,69^, and congenital heart disease^70^, our work provides insight into how WDR62-dependent mechanisms in MT organization may contribute to disease beyond the germline.

## MATERIALS AND METHODS

### Fly stocks and genetics

All fly stocks were grown on standard medium at room temperature. The following stocks were used in this study: Wdr62-GFP (II, from Bloomington *Drosophila* Stock Centre (BDSC) #66372), mataα4-Gal-VP16-V37 (III, BDSC#7063), mataα4-Gal-VP16-V2H (II, BDSC#7062), patronin^EY05252^ (II, BDSC#16644), Patronin-mCherry (II, gift from Uri Abdu, Department of Life Sciences, Ben-Gurion University of the Negev, Israel), Patronin-YFP (II) and Staufen-RFP (III) (gift from Dr. Daniel St Johnston, The Gurdon Institute and the Department of Genetics, Cambridge, UK, (Nashchekin et al., 2021; Nashchekin et al.,2016), Jupiter-GFP and Jupiter-mCherry (III, gift from Dr. Mónica Bettencourt-Dias, Instituto Gulbenkian de Ciência (IGC), Oeiras, Portugal), UASp-EB1-GFP (II, gift from Dr. Antoine Guichet, CNRS, Institut Jacques Monod, Paris, France). The RNAi lines used were:P{TRiP.GL01286}attP2 (III, BDSC #41858) for Short-Stop (Shot) and P{TRiP.GLC01394}attP2 for Wdr62 (III, BDSC #53242). Both lines are GL RNAi lines constructed using the VALIUM22 vector, which drives expression in the female germline. Knockdown for Katanin with UAS-RNAi was carried out with the line P{TRiP.HMC06296}attP40 (II, BDSC#66000). This line is a HM RNAi line constructed using the VALIUM20 vector, which drives expression in the female germline. To deplete these proteins, these RNAi lines were crossed with mataα4-Gal-VP16-V37 driver line. A UAS-RNAi line for mCherry was used as control (III, BDSC#35785).

The following combination of fly genotypes were generated for this study:

V32-Gal4; Staufen-RFP

V32-Gal4; EB1-GFP

V32-Gal4; Jupiter-GFP

Wdr62-GFP; Jupiter-mCherry

Wdr62-GFP; mCherry RNAi

Wdr62-GFP; Shot RNAi

Wdr62-GFP; Wdr62 RNAi

Patronin-YFP; mCherry RNAi

Patronin-YFP; Shot RNAi

Patronin-YFP; Wdr62 RNAi

V32-Gal4 was recombined to Jupiter-GFP generating flies V32-Gal4::Jupiter-GFP (III). Patronin^05252^; V32-Gal4::Jupiter-GFP

### Egg laying and hatching

Groups of 7 day-old well fed females with 5 males were anaesthetized using CO2 exposure and transferred to apple juice agar plates, where they were allowed to lay eggs for 24h. Flies were then removed, and the number of eggs was assessed. After 48h at 25°C, the eggs hatched were evaluated. Egg laying was calculated by dividing the number of eggs by the number of living females.

### Antibodies used

The following primary antibodies were used: mouse anti-Shot (1:20; Hybridoma), mouse anti-gurken (1:20; Hybridoma), chicken anti-D-plp (1:400, Sigma), rat anti-α-tubulin (1:500, Santa Cruz Biotechnology). Secondaryantibodies were from ThermoFisher.

### Ovaries immunostaining

#### For mid-stages

Ovaries were dissected from 3-4 day-old well fed females. The protocol was adapted from Januschke et al., 2006. For ovary dissection, females were anesthetized with ether and transferred to pre-warmed (25°C) BRB80 buffer (80mM Pipes pH 6.8, 1mM MgCl2, 1 mM EGTA) supplemented with 1× protease inhibitor (5056489001, Roche). Ovaries were extracted with pre-cleaned forceps. Individualized ovaries were then incubated for 1 hour at room temperature (RT) in BRB80 with 1% Triton X-100 without agitation, followed by a 15 minutes fixation step at –20°C in pre-cooled methanol. 3 wash steps of 5 min each and overnight permeabilization were done in PBST (1× PBS with 0.1% Tween) at 4°C. Blocking for 1h was done in PBST with 2% BSA (Sigma #A2153). Primary antibodies were incubated overnight at 4°C in PBS with 1% BSA (Sigma #A2153) (PBSB) with agitation, followed by 3 wash steps of 15 min each. Secondary antibodies were diluted in PBSB and incubated for 2h at RT. Ovaries were washed 2 times for 15 min in PBT and 2 times for 15 min in PBS, and DNA was stained using DAPI (Vector Laboratories).

Alternatively, to visualize Gurken and Staufen-RFP proteins (**Fig. 2**), a modified fixation protocol was used. Individualized ovaries were dissected and fixed for 1h without agitation, in a solution mix containing 1XPEM (80mM PIPES + 2mM EGTA + 1mM MgCl2), 1% Triton-X-100 and 4% paraformaldehyde. 3 wash steps of 15 min each were done in PBS with 0,3% Triton-X-100 (PBS-Tx). Ovaries were blocked for 1h in PBS, 0,05% Tween-20 with 2% BSA (Sigma #A2153). Primary antibodies were incubated overnight at RT in PBS, 0,05% Tween-20 with 1% BSA (Sigma #A2153) (PBT1) with agitation, followed by 3 wash steps of 15 min each in PBS-Tx. Secondary antibodies were diluted in PBT1 and incubated for 2h30min at RT. Ovaries were washed 3 times for 10 min in PBS with 0,05% Tween-20. DNA was stained using DAPI (Vector Laboratories). The coverslips were mounted using the same mounting medium (Dako Omnis) for both fixation methods. Images were acquired on a Zeiss LSM980 system, using a 63x 1.4NA Oil immersion objective with ZEN Blue 3.3. Serial sections were acquired every 19 µm.

#### For meiosis

Ovaries were dissected from females fed on wet yeast 72 hours into BRB-80 buffer (80mM Pipes pH 6.8, 1mM MgCl2, 1 mM EGTA) supplemented with 1× protease inhibitor. Ovaries were extracted with pre-cleaned forceps. Individualized ovaries were then incubated for 1 hour at 25 °C in BRB80 with 1% Triton X-100 without agitation, followed by a 15 minutes fixation step in 500 μL 4% paraformaldehyde stabilized with PBS. During the last 5 minutes of fixation, up and down was done with a micropipette to gently separate the oocytes. Oocytes were then washed for 10 minutes, three times in PBS with 0.1% Triton X-100 and once with PBS. Oocytes were mounted on a glass slide in Vectashield with DAPI (H-1200, Vector Laboratories). For high resolution imaging of the spindle, images were acquired as Z-series on a Zeiss LSM980 system, using a 63x 1.4NA Oil immersion objective with ZEN Blue 3.3. All images were acquired with the same exposure.

### Live imaging of *Drosophila* Egg Chambers

Stage 7–8 oocytes were dissected directly in Voltalef 10S oil (VWR Chemicals) and mounted on glass-bottom dishes (MatTek) for live imaging. EB1–GFP imaging was performed using a Zeiss LSM 980 confocal microscope equipped with an Airyscan 2 detector and a 63×/1.40 NA oil immersion objective. GFP was excited using a 488 nm laser line. Time-lapse movies were acquired as single optical sections (one z-plane) for at least 4 minutes with a frame interval of 500 ms (2 frames per second) to enable tracking of individual microtubule (MT) plus-end comets. Image acquisition and Airyscan processing were performed using ZEN software (Zeiss). Jupiter–GFP imaging was performed using a Zeiss Cell Observer SD spinning disk confocal system equipped with a CSU-X1 scanner and a 63×/1.40 NA oil immersion objective, controlled with ZEN Blue 3.4 software (Zeiss). GFP was excited using a 488 nm laser line. Time-lapse movies were acquired as z-stacks of 10 optical sections every 2.23 s for at least 5 minutes.

### Quantification of Eb1-GFP dynamics

The quantification of Eb1-GFP dynamics was performed based on the protocol described by Cabrita et al. 2026 (*J.Vis.Exp, in press*), with specific modifications to accommodate the low signal-to-noise ratio observed in single and double mutant lines. Additional filtering was added to the protocol to avoid having false negatives and false positives. During the spot detection phase (step 3 of the protocol), the spot size was set to §0.4 µm and the quality threshold was reduced to 1.0 and a otsu auto-thresholding method was applied to the distribution of intensities of the spots detected. Particle tracking was executed using the Advanced Kalman Tracking algorithm with a search radius of 0.3 µm and feature weights set to 1 for both spot quality and Channel 3 intensity. In the final calculation step (Step 4 of the protocol) we used a value of 5 to remove tracks with fewer than 5 frames and removed tracks with less then 0.3 angular consistency.

### Quantification of Microtubule density in live oocytes

For quantification of microtubule density, Jupiter-GFP time-lapse movies were processed in Fiji (ImageJ). Z-stacks (10 optical sections) were first projected using maximum intensity projection. The resulting time series (135 timepoints) was then averaged across time to generate a single representative image per oocyte. Stage 7–8 oocytes were manually segmented using the polygon selection tool, and the mean fluorescence intensity within the oocyte area was measured and used as a proxy for microtubule density. All samples were imaged using identical acquisition settings. Quantification was performed using Fiji (ImageJ).

### Quantification of Gurken staining at the antero-dorsal corner of the oocyte

Gurken fluorescence was quantified using Fiji (Image J). For each stage 8 oocyte, Z-slices containing detectable Gurken signal were selected. Background fluorescence was estimated by measuring the intensity of three different regions lacking specific signal, and the mean value was subtracted from the original image. Following background subtraction, Gurken-positive region was defined by manually drawing a region of interest encompassing the entire signal. Total fluorescence intensity was then measured on a sum Z-projection image.

### Quantification of Staufen–RFP at the posterior cortex

Stage 10 oocytes expressing Staufen-RFP were analyzed using Fiji (ImageJ). For each oocyte, a sum Z-projection was generated from the slices containing detectable Staufen signal. Background fluorescence was measured from three regions outside the oocyte and the mean value was subtracted from the images. The Staufen-positive region at the posterior cortex was defined using an intensity threshold, and the total fluorescence intensity within this region was measured.

### Quantification of Shot and Wdr62 intensity in the oocyte

The mean fluorescence intensity of α-Shot and Wdr62::GFP was measured in stage 9-10 immunostained oocytes using Fiji (Image J). To assess cortical enrichment, a segmented line selection tool with a width of 20 was used along the antero-lateral cortex. Cytoplasmic enrichment was measured by manually segmenting the oocyte using the polygon selection tool and quantifying the mean fluorescence intensity within this region. Background fluorescence was measured outside the oocyte and subtracted from all intensity measurements.

### Qualitative analysis of protein localization (Staufen, Gurken and Patronin)

Staufen-RFP, Gurken and Patronin-YFP localization was assessed qualitatively by classifying oocytes as displaying either normal or abnormal localization patterns. Normal localization was defined based on the distribution in control oocytes (posterior enrichment for Staufen, dorsal-anterior localization for Gurken, and cortical localization for Patronin-YFP), whereas deviations from these patterns were classified as abnormal localization. Percentages represent the proportion of oocytes in each category.

### Qualitative analysis of Wdr::GFP co-localization with D-plp

Wdr62::GFP co-localization with the centriole marker, D-plp, was assessed qualitatively in oocytes from the germarium, stage 4-5, and stage 9 egg chambers. Circular regions were defined around D-plp signal using the oval selection tool in FIJI (Image J). The spatial relationship between Wdr62::GFP and D-plp signal was then visually evaluated. Oocytes were classified into three categories: co-localized (overlapping signals), non-colocalized (no overlap), or adjacent (Wdr62::GFP signal positioned immediately next to, but not overlapping, D-plp). Percentages represent the proportion of oocytes in each category.

### Quantification of meiotic spindle

A minimum of 10 oocytes at stage 14 were analyzed for each biological replicate and each condition. Spindle presence, morphology, length and width were analyzed for spindle microtubules. Spindle morphology was assessed qualitatively according to spindle bipolarity, microtubule organization and overall spindle morphology. Based on this assessment, spindles were categorized as normal or abnormal. Percentages represent the proportion of oocytes in each category. Spindle length and width were measured using Fiji (Image J) line tool, by drawing a line from one spindle pole to the other, for length, and a line across the widest part of the spindle, for width.

### Quantification of the dorsal appendage morphology

Groups of 7 day-old well fed females with 5 males were anaesthetized using CO2, and transferred to apple juice agar plates, where they were allowed to lay eggs for 24h. Flies were then removed, and the dorsal appendages of the collected eggs were assessed. Dorsal appendages were classified into five categories: symmetrical, fused, short, absent, and unilateral. Percentages represent the proportion of oocytes exhibiting either normal (symmetrical) or abnomal phenotypes (fused, short, absent, and unilateral).

### Determining the timing of NEBD

Females were collected and fed on wet yeast for 48 hours. Ovaries were dissected in halocarbon oil 700 (H8898, Sigma Aldrich) on a slide, and egg chambers were carefully separated. The oocytes were immediately imaged on a Zeiss SteREO V8. The presence or absence of a nucleus on late stage 13 oocytes and the number of stage 14 oocytes were assessed.

### Statistical analysis

All experiments were performed with at least three independent biological replicates per condition. The number of cells and egg chambers analysed for each experiment are detailed in the respective figure legends.

#### Quantification of Eb1-GFP dynamics

Differences between experimental conditions (for >2 groups) were assessed using one-way ANOVA followed by the Tukey’s multiple comparisons test, Games–Howell multiple-comparisons test, or the Kruskal–Wallis test with multiple comparisons, as appropriate. Directional preferences within the same ROI were assessed using the Friedman test (global comparison), followed by the Wilcoxon Signed-Rank test (for pairwise comparisons).

#### Quantification of Gurken, Staufen and microtubule (Jupiter-GFP) intensity in the oocyte

To quantify the impact and (statistical) significance of each RNAi on Gurken and Staufen localization, and Microtubule density, independent differences between experimental conditions were assessed using the Kruskal–Wallis test with multiple comparisons (for >2 groups).

#### Quantification of Shot and Wdr62 intensity in the oocyte

To quantify the impact and (statistical) significance of each RNAi, independent differences between experimental conditions were assessed using the Mann-Whitney U test unpaired (for two groups).

#### Qualitative analysis of protein localization (Staufen, Gurken and Patronin)

To quantify the impact of each RNAi and (statistical) significance of each RNAi on Gurken, Staufen and Patronin localization, independent differences between experimental conditions were assessed using the Fisher’s exact test.

#### Egg laying and hatching

To quantify the impact of each RNAi on egg laying and hatching, Kruskal-Wallis multiple comparison test was performed between the conditions.

#### Quantification of the dorsal appendage morphology

To quantify the impact of each RNAi and (statistical) significance of each RNAi on the dorsal appendage morphology, a Chi-square test was performed between conditions.

#### Quantification of meiotic spindle

To test the statistical significance of each RNAi on spindle morphology, a Chi-square test was performed between conditions. For spindle length and width, a t-test was used in Wdr62 depletion experiments and Kruskal-Wallis multiple comparison test for the experiments with Wdr62 and Patronin depletion.

#### Determining the timing of NEBD

To quantify the impact of each RNAi on the timing of NEBD, the percentage of stage 13 oocytes with NEBD and without NEBD were compared between conditions using Chi-square test. To assess the impact on the number of stage 14 oocytes, a t-test was performed between conditions.

## ACKNOWLEDGMENTS

We are very grateful to Zita Carvalho-Santos (Gulbenkian Institute for Molecular Medicine, Portugal), Mónica Bettencourt-Dias (Centre for Genomic Regulation, Spain) and Carlos Conde (Instituto de Investigação e Inovaçãoe em Saúde, Portugal) for insightful discussions and critical reading of the manuscript. We thank Antoine Guichet (Institut Jacques Monod, France) for the UASp-EB1-GFP fly line, to Daniel St Johnston (Gurdon Institute, University of Cambridge, United Kingdom) for the Patronin-YFP fly line, and to Uri Abdu (Ben-Gurion University of Negev, Israel) for generously sharing with us the UASp-Patronin-mCherry line; V32-Gal4 fly line. We also thank Clemens Cabernard (University of Washington, USA) for sharing fly lines. We also acknowledge the technical support and assistance of the Microscopy Facility at NOVA Medical School, supported by PPBI (POCI-01-0145-FEDER-022122), and the Fly Facility at NOVA Medical School, supported by CONGENTO (LISBOA-01-0145-FEDER-022170). This work was supported by funding awarded to A.P.M. (2024/158225/PEX), by the Research Unit iNOVA4Health – Programa de Medicina Translacional (UID/04462/2025), and by the Associated Laboratory LS4FUTURE (LA/P/0087/2020), all financially supported by the Fundação Para a Ciência e Tecnologia (FCT) / Ministério da Educação, Ciência e Inovação. A.P.M. is supported by an FCT researcher contract under the CEECInd programme (CEECIND/02842/2020), and J.C. is supported by an FCT PhD Fellowship (2023/03665/BD).

## Notes

### Competing Interest Statement

The authors have declared no competing interest.

